# Membrane curvature during membrane rupture and formation of pentagonal pyramidal superassemblies by a pore-forming toxin, *Vibrio cholerae* Cytolysin, using single particle cryo-EM

**DOI:** 10.1101/2025.07.19.665663

**Authors:** Suman Mishra, Kausik Chattopadhyay, Somnath Dutta

## Abstract

In this cryo-electron microscopy study, we provide mechanistic insights into how an archetypical β-barrel pore-forming toxin (β-PFT), *Vibrio cholerae* Cytolysin (VCC), ruptures the membrane lipid bilayer by inducing membrane curvature. We demonstrate how VCC oligomers cluster together and drastically increase local membrane curvature, thereby causing membrane blebbing. In addition, we also show how these PFTs after rupturing the host membrane tend to form symmetric super-molecular assemblies to stabilize their hydrophobic transmembrane rim domains. We further provide another example of membrane rupture with gamma hemolysin, a Staphylococcal bicomponent β-PFT. These insights will usher new studies on membrane curvature due to protein crowding and broaden our mechanistic understanding of how this largest class of bacterial protein toxins induce host cellular death.

## INTRODUCTION

Pore-forming toxins (PFTs) constitute the largest class of bacterial protein toxins(1). Secreted as water-soluble monomers, they often undergo a dramatic conformational change upon selectively binding to specific lipid/protein receptors on host cell membranes-forming pores that induce osmotic imbalance and cellular death. Initially, the monomeric toxin binds to the lipid membrane, where it undergoes a conformational change. This change causes the pre-stem region to separate from the core region of the protein. The pre-stem region then inserts itself into the membrane, while the core region remains largely intact (**Figure 1*A-C***). During this process, the pre-stem is able to adopt a transmembrane beta-barrel structure that penetrates the lipid bilayer and punches the holes in the membrane. These beta-barrel-mediated holes behave like a channel, which allows the toxin to extract ions and nutrients from the host cells and disrupt membrane integrity. Traditionally, cellular death induced by such PFTs was attributed to their channel-like activity. However, in the recent past, several studies on a wide repertoire of PFTs have shown that these toxins destroy host cellular integrity by also interacting with surface proteins and inducing downstream cell signaling(1). Additionally, PFTs serve as an archetype of membrane-interacting proteins and some studies have demonstrated how PFTs are influenced by membrane properties and vice versa(2–8). One interesting phenomenon has been consistently observed for several unrelated PFTs in membrane environment, that they oligomerize and rapidly aggregate on the lipid membrane, limiting the application of near-native model membranes such as liposomes(6, 9, 10). However, one recent structural study also showed how MakA toxin from *Vibrio cholerae* formed supramolecular filamentous structures with liposomes, illustrating how structural studies significantly depend on the type of model membrane used(11). Furthermore, we also previously demonstrated with a Staphylococcal PFT, γ-hemolysin, that the monomeric subunits oligomerize upon liposomes and form clusters and supramolecular assemblies after membrane rupturing(6). However, a direct visualization of the impact of such PFT oligomers on the lipid membrane properties was missing thus far. In this study, we provide direct evidence of membrane curvature induced by clustering of an archetypical β-PFT, *Vibrio cholerae* Cytolysin (VCC). This study proposes the idea that membrane damage caused by PFTs can also be attributed to rupturing due to protein clustering in addition to channel-like activity. We further demonstrate how VCC heptamers after rupturing lipid membranes tend to form supramolecular assemblies and acquire a rather unusual pentagonal pyramidal geometry. This, along with our previous study on γ-hemolysin, also demonstrates the possible application of these different pore-forming toxins to form nanocages/scaffolds of desired geometry.

**Figure 1.**
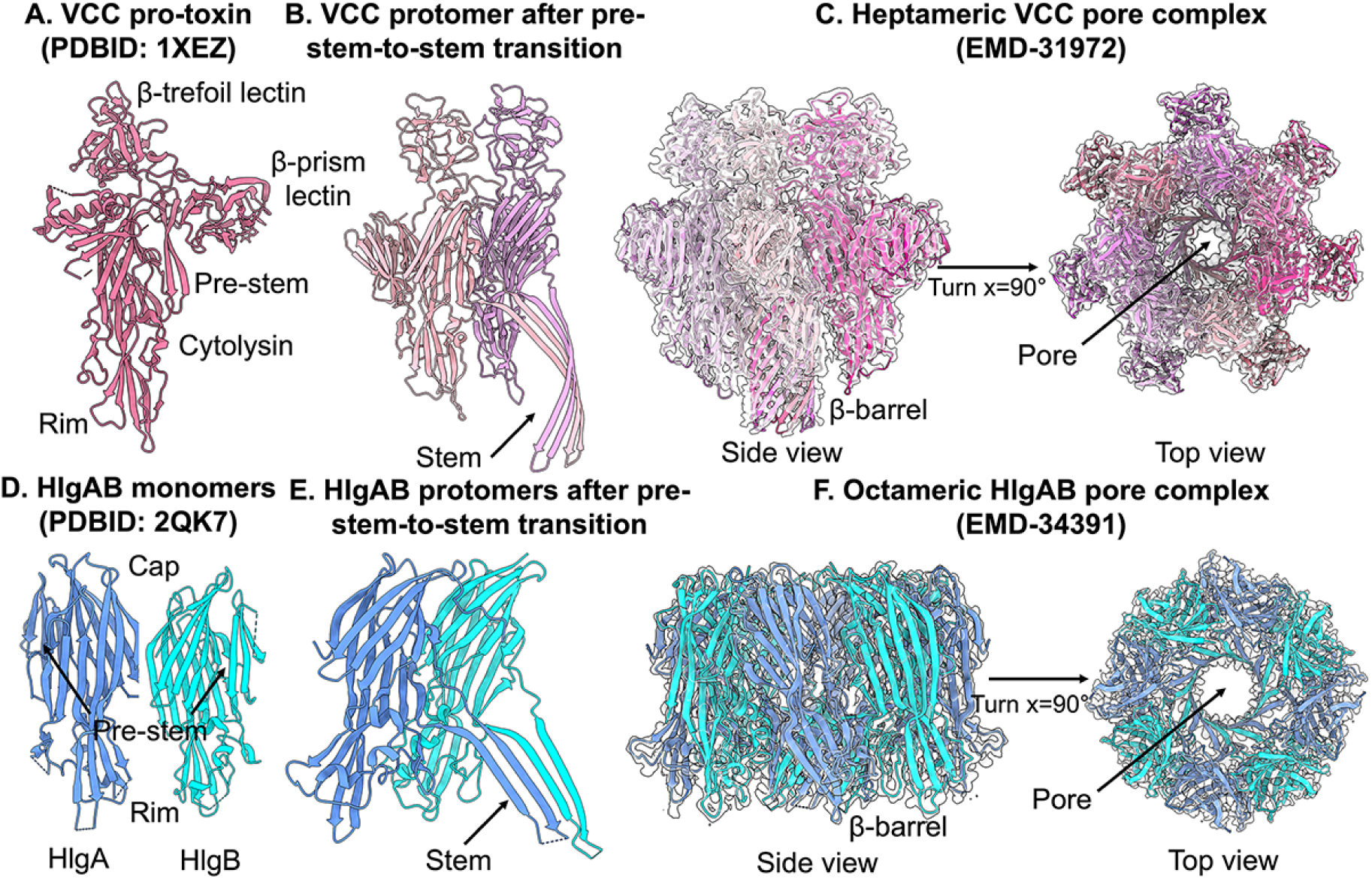
*Vibrio cholerae* Cytolysin and γ-hemolysin, a bicomponent PFT from *S. aureus*. (A) Monomeric crystal structure representing the pro-toxin form of VCC (PDBID: 1XEZ). This pro-toxin undergoes proteolytic cleavage that permits oligomerization of toxin monomers on host cholesterol-rich membranes. (B) Insertion of hydrophobic pre-stem domain of VCC monomer in the lipid membrane. (C) Cryo-EM map of heptameric VCC pore with fitted atomic model (EMD-31972). The water-soluble monomers undergo large conformational changes to form an amphipathic transmembrane structure with a β-barrel. (D) Monomeric structures of the two toxin subunits of γ-hemolysin-HlgA and HlgB (PDBID: 2QK7). (E) Insertion of hydrophobic pre-stem domain of HlgA and HlgB monomers in the lipid membrane. (F) Cryo-EM structure of octameric HlgAB in a lipidic environment (EMD-34391). The two toxin subunits individually cannot oligomerize to form a functional pore on the host membrane, but together they form an octameric pore complex with alternating HlgA and HlgB subunits.

In this current study, we employed single particle cryo-EM analysis to understand the membrane rupturing phenomenon by VCC. During membrane rupture, membrane curvature should be altered by PFTs. Using our cryo-EM 2D classification strategy, we were able capture several lipid curvature changes induced by VCC. Furthermore, we were able to capture a super molecular assembly of VCC in the presence of a large lipid vesicle.

## RESULTS

### Visualizing the membrane curvature in the presence of VCC

We considered large unilamellar vesicles constituted of asolectin and cholesterol as an appropriate model membrane to study the impact of VCC oligomers on the lipid architecture. The large membrane surface provided by liposomes closely mimics the host cellular membranes and permits studying clustering of proteins, providing a significant advantage over other model membranes such as lipid nanodiscs and detergents. We previously provided the high-resolution structure of VCC in a lipid environment(5). So, for this study, we focused on previously unexplored aspects of membrane curvature induced by PFTs. We acquired a cryo-EM dataset of ∼3500 micrographs of VCC-embedded liposomes using a 200 kV Talos Arctica to employ single particle analysis.

Cryo-EM dataset of VCC with lipid membrane was highly heterogeneous with presence of different orders of protein clustering on lipid membrane and also supramolecular lipid-protein assemblies outside the liposomes (Figure 2). We examined several cryo-EM images of VCC-embedded liposomes, which showed the various lipid curvature during membrane rupture. Furthermore, we also observed that membrane curvature varies in the presence of toxin molecules. However, this was an extremely challenging task to observe the membrane curvature phenomenon and count the number of protein molecules bound with liposomes from the noisy raw cryo-EM micrographs. Therefore, we performed several rounds of 2D classification targeting different assemblies (Figure 3). To accommodate clusters of VCC oligomers on the lipid membrane, we chose to extract VCC particles embedded in liposomes using a large box size of 480 pixels (1.17 Angstroms/pixel). Several rounds of 2D classification strikingly revealed well-aligned 2D class averages with different numbers of VCC oligomers embedded in the membrane (Figure 3). Importantly, we saw that VCC oligomers induced membrane curvature (Figure 4). Liposomes inherently have a curved surface, and we measured the same in a 2D class devoid of proteins to be ∼172 degrees. When a single VCC oligomer formed a pore and punctured the membrane bilayer, this resulted in an increase in membrane curvature and a decrease in the measured angle, which was ∼148 degrees (Figure 4). Interestingly, when two VCC heptamers clustered together on the membrane surface, this resulted in a dramatic increase in membrane curvature with the measured angle being 139 and 127 degrees for two different 2D classes representing different states of VCC clustering on the membrane (Figure 4). Furthermore, we observed the presence of three VCC oligomers clustering together, where they can be seen inducing the maximum curvature of 105 degrees and pinching off a stretch of the lipid membrane (Figure 4). The different 2D classes obtained in this study and their corresponding state of membrane curvature along with sample size is provided in Table 1. It is important to note here that in cryo-EM single particle analysis, all the particles corresponding to a particular 2D class average might not necessarily correspond to the same state and might also include junk particles or particles from a different state. This makes it difficult to comment on relative stability of different membrane-bound states. In addition, a few classes are represented by a relatively small population of particles due to high heterogeneity present in the dataset which does not exclude the possibility that some good particles might also be lost in junk classes during previous rounds of 2D classification.

**Figure 2.**
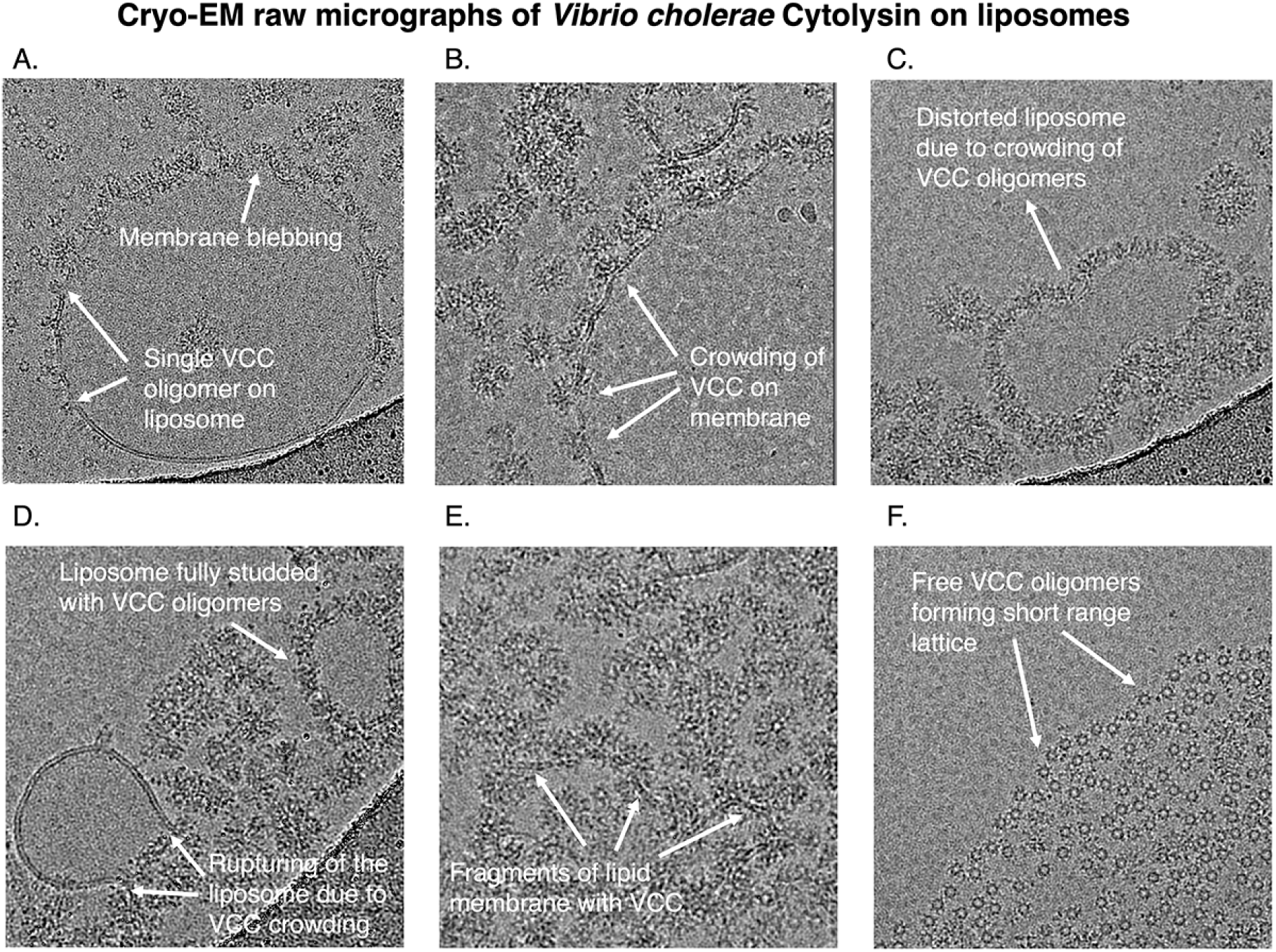
Cryo-EM micrographs illustrating membrane curvature induced by VCC. Representative raw micrographs of VCC-embedded liposomes illustrating different states of membrane rupture. Representative white arrows represent sites where VCC induces membrane curvature in the liposomes. (A) An intact liposome decorated with transmembrane ring-like VCC oligomers. Single oligomers and clusters of several oligomers are visible on the same vesicle. (B) Several clusters inducing membrane blebbing are visible in the raw micrographs. These vesicles otherwise appear to retain their globular architecture. (C) Distorted dumbbell proteo-liposome architecture can be observed in this micrograph. (D) A ruptured liposome can be seen with the rupture occurring at the site of VCC clusters. (E) Small fragments of liposomes can be seen with tethered VCC oligomers. (F) VCC oligomers can be seen outside the lipid membrane forming a small-range lattice.

**Figure 3.**
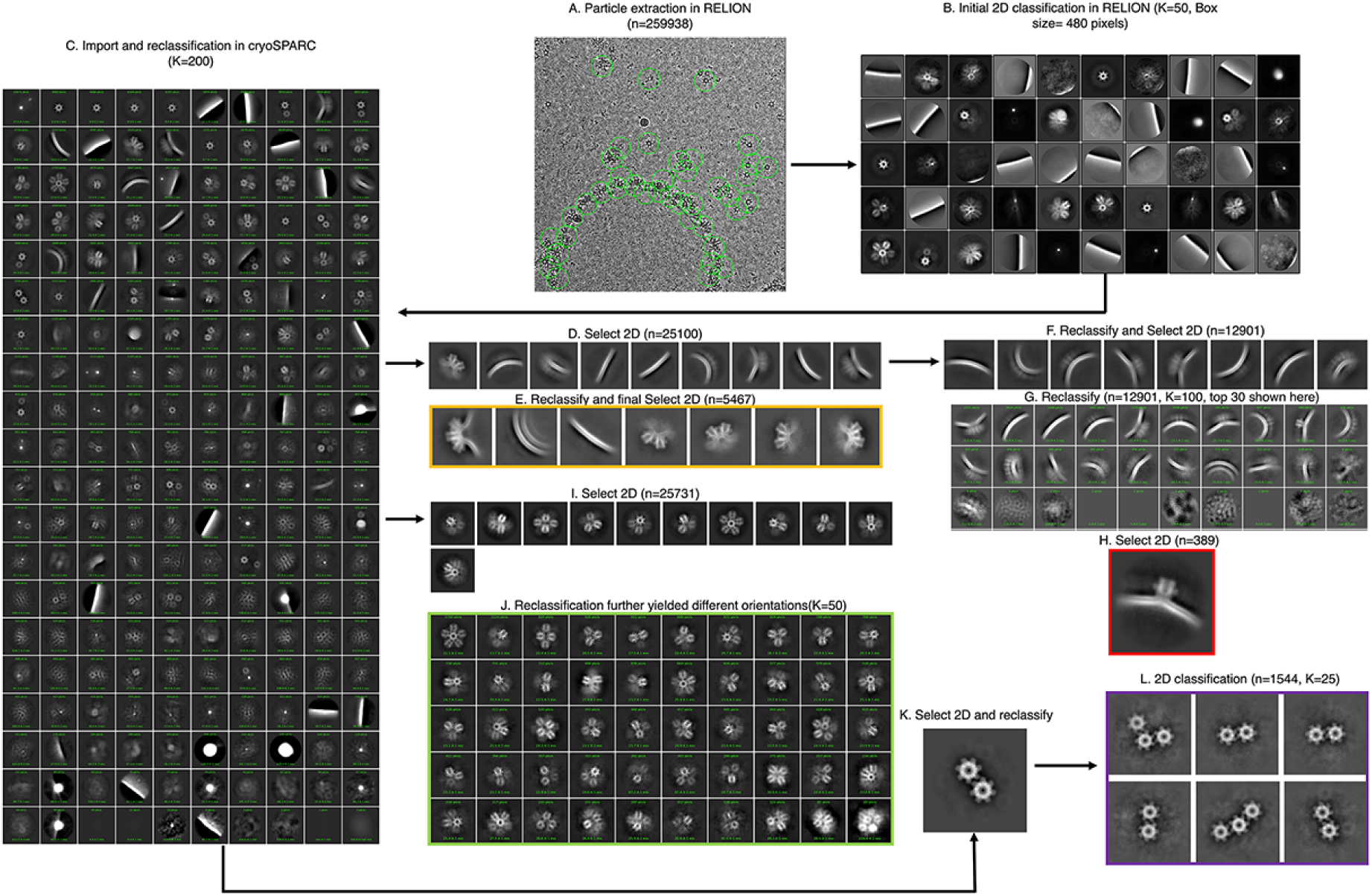
Data Processing pipeline for cryo-EM study of VCC-embedded liposomes. A rigorous 2D classification approach upon extracting with a large box size of 480 pixels permitted us to obtain 2D class averages corresponding to different states of VCC such as a single oligomer on the liposome membrane, multiple oligomers inducing different degrees of membrane curvature, pentagonal pyramidal supramolecular assemblies and several other lipid-protein complexes. Picking and extraction were performed in RELION3.1 while 2D classifications were performed in cryoSPARCv4.1.

**Figure 4.**
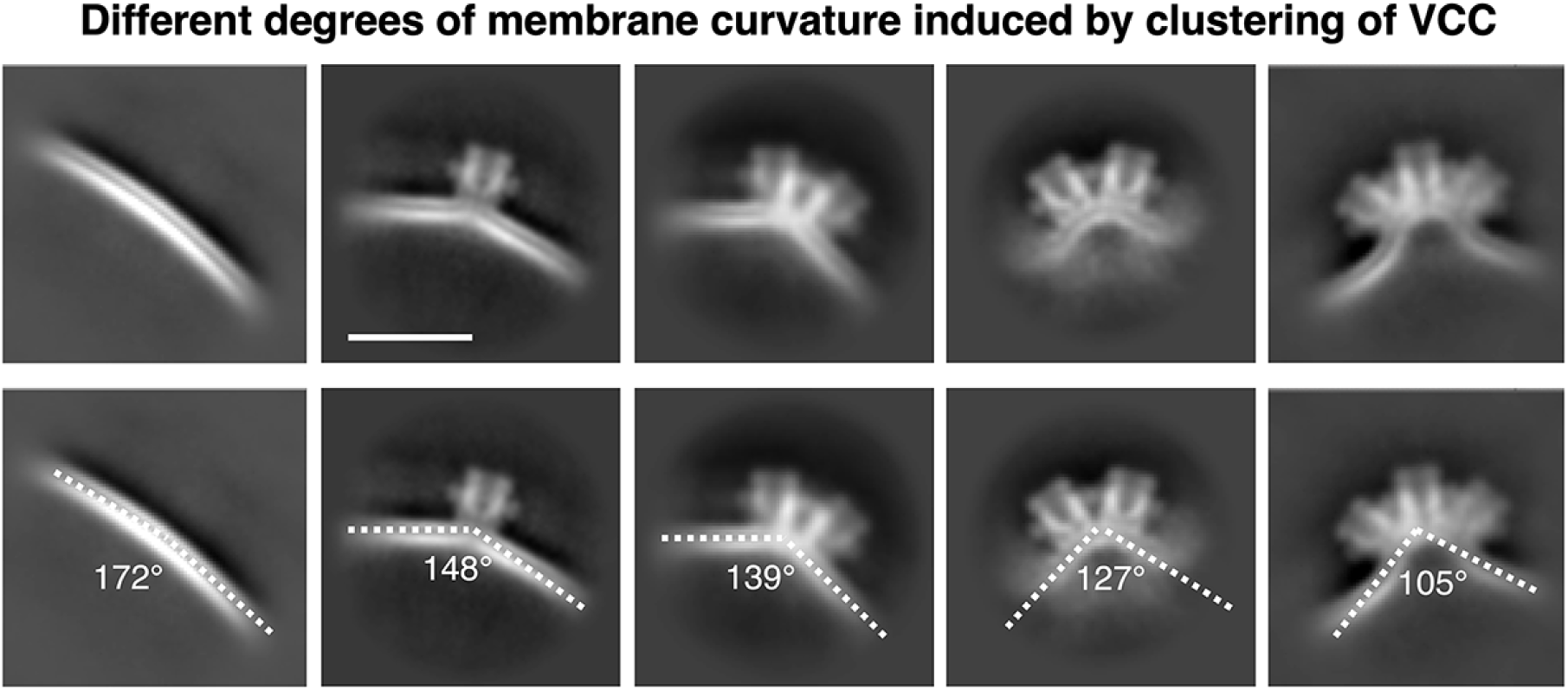
Membrane curvature induced by clustering of VCC oligomers. Representative cryo-EM 2D class averages illustrate different degrees of membrane curvature induced by clustering of VCC oligomers. Top row and bottom row comprise raw 2D classes and 2D classes with marked curvature respectively. Calculations to measure the angles of curvature were made using an online tool, (https://www.ginifab.com/feeds/angle_measurement/)

**Table 1:** Different states obtained in 2D classification and corresponding sample size.

| Class description | Impact on membrane | Number of particles |
| --- | --- | --- |
| Only liposomes, no toxin | 172 degrees local curvature | 3747 |
| One oligomer adhered to membrane | 148 degrees bent membrane | 389 |
| Two oligomers clustering together | 139-127 degrees bent membrane | 3560 |
| Three oligomers clustering together | 105 degrees bent membrane | 2151 |
| VCC lattice | post membrane rupture state | 1544 |
| VCC-lipid fragments | post membrane rupture state | 306 |
| Pentagonal pyramidal assemblies | post membrane rupture state | 25731 |

Our cryo-EM data provides strong evidence for membrane curvature induced by protein clustering(12), and also illustrates at near atomic resolution different states of how VCC oligomerization influences host membrane integrity. Our recent data also suggest that the bi-component toxin γ-hemolysin also followed the same membrane blebbing during membrane rupture and formed lipid-protein clusters (Figure 6).

### Super-molecular assembly of VCC with ruptured lipid membrane

We previously demonstrated with Staphylococcal bicomponent γ-hemolysin that its oligomers tended to form supramolecular assemblies after membrane rupture in both liposomes as well as erythrocytes(6). For γ-hemolysin, the superassembly consisted of an octahedral arrangement of six octamers comprising alternate arrangement of the two toxin components HlgA and HlgB. The octahedral superassembly that comprised 48 toxin monomers was stabilized primarily by hydrophobic residues in the rim domain that were known to be crucial for membrane interaction.

In our current study with VCC, we further investigated the possibilities of VCC heptamers to form larger assemblies for stabilization of the rim domain after the membrane rupture. We further picked and extracted VCC particles that were outside the liposomes in our cryo-EM dataset with a large box size of 480 pixels and subjected them to 2D classification (Figures 2 and 3). Interestingly, we could identify not only different orientations of an ordered superassembly but also other lipid-protein clusters that illuminated the fate of these oligomers after rupturing of lipid membranes (Figure 5A-B). Upon generating a 3D reconstruction from the different orientations and fitting the atomic model of VCC, we observed that six subunits of VCC heptamers were arranged in pentagonal pyramidal geometry which has rarely been previously observed for any other proteins (Figure 5C).

**Figure 5.**
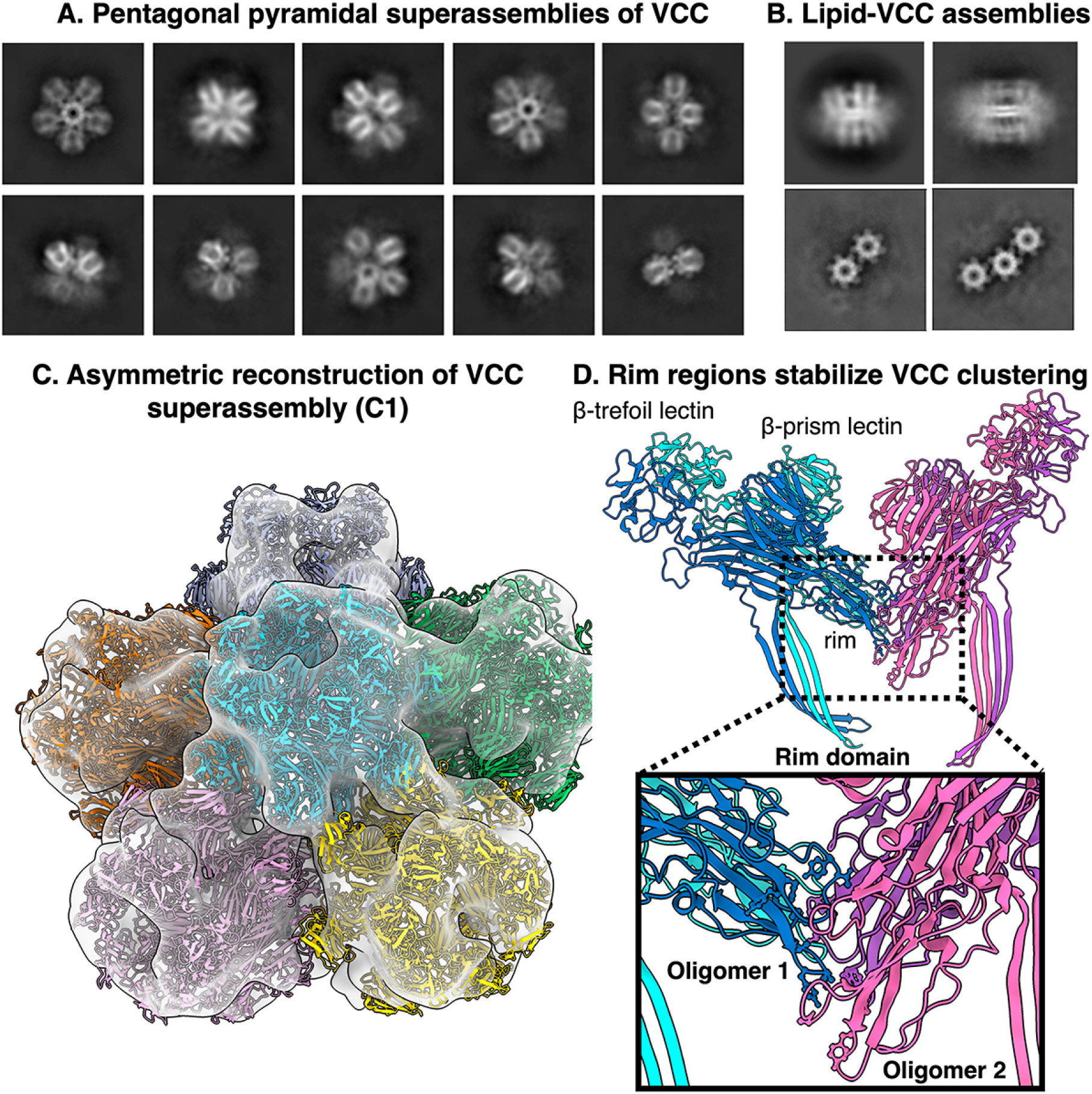
Pentagonal pyramidal supramolecular assembly of VCC heptamers. (A) 2D class averages corresponding to different orientations of these superassemblies. (B) 2D class averages illustrating other lipid-protein clusters. (C) Cryo-EM map of VCC pentagonal pyramidal superassemblies with fitted atomic model of heptameric VCC in six different colors. Molecular interactions between two heptamers, cyan and pink, are shown in (D). The rim domain of monomeric VCC from one heptamer is sandwiched between two monomers from the neighboring heptamer. Oligomers 1 and 2 are indicated with different colors.

**Figure 6.**
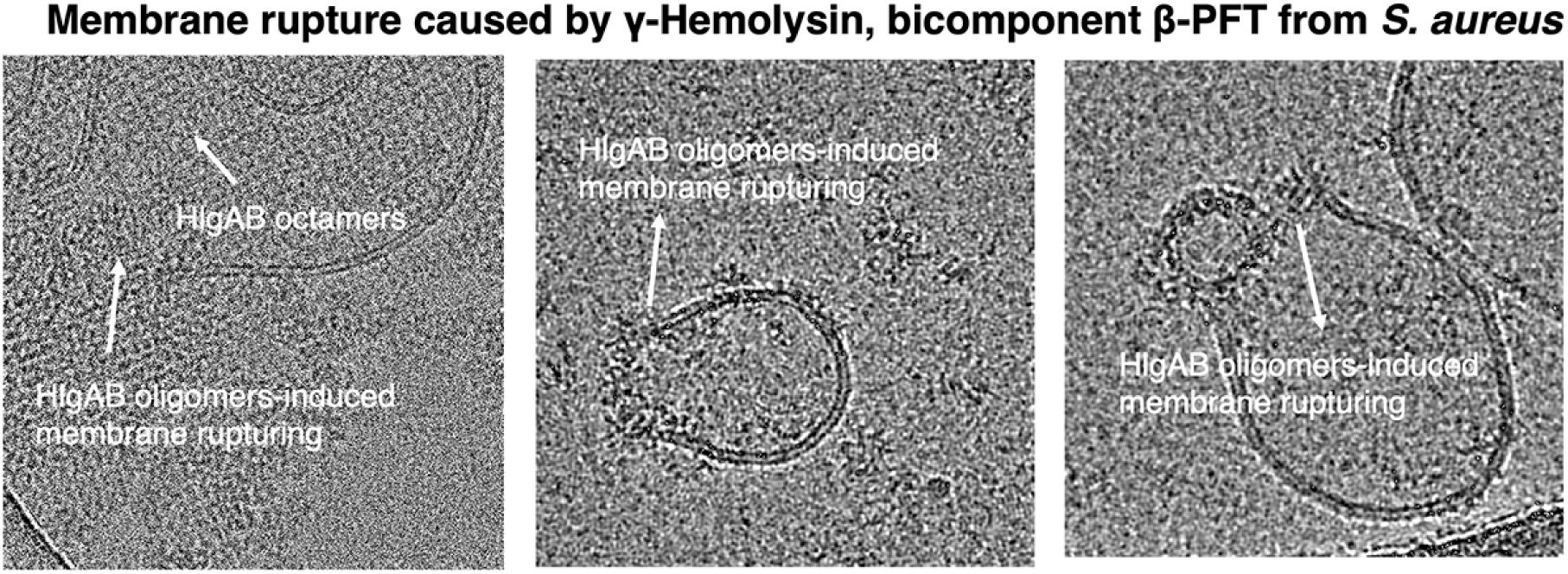
Membrane rupturing by a bicomponent PFT γ-hemolysin from *Staphylococcus aureus*. Representative cryo-EM raw micrographs of Staphylococcal γ-hemolysin with phosphatidylcholine-cholesterol liposomes illustrating membrane damage due to γ-hemolysin oligomers.

At the outset, the pentagonal pyramidal architecture permits the hydrophobic rim regions of each VCC heptamer to be packed against each other, thus stabilizing the structure similar to the octahedral assembly of γ-hemolysin. Since the atomic model fitted well in our cryo-EM map, we wanted to further pinpoint the residues for VCC that stabilized its superassembly (Figure 5C-D). In our previous high resolution cryo-EM map of VCC in lipid membrane, we could identify that the rim region contributes to lipid binding through residues Y235-T236, T237-L238, Y241-F242, L361-W362-V363, Y417, H419-Y420-Y421, V422-V423 and H426. In the pentagonal assembly, we observe the loop comprising Y235-Y241 from a protomer (say Protomer A) from first heptamer sandwiched between the most extended loop Y421-V423 of one protomer (Protomer B) and L361-V363 loop of another neighboring protomer (Protomer C) from the second heptamer (Figure 5D). However, from our low resolution cryo-EM map accurate distances between these interacting loops were not calculated (Figure 5C-D).

Nevertheless, the interaction of these loops from the rim domain support our previous understanding of the lipid interacting residues. Our observations with VCC pentagonal pyramidal superassemblies are concomitant to those with γ-hemolysin octahedral supramolecular assemblies and confirm that multiple PFTs have the tendency to rupture the lipid membranes and form larger clusters unlike what we traditionally observe for native channel-like proteins.

## DISCUSSIONS

In this study, the use of large unilamellar vesicles (LUVs) as a model membrane was pivotal since it enabled us to directly visualize the clustering of VCC oligomers and its effect on membrane curvature. Liposomes closely mimic the bilayer architecture of native membranes, thus providing us with a near-physiological platform to study lipid-protein interactions. Unlike simpler model membrane systems such as detergent micelles and lipid nanodiscs, these liposomes offer a large surface area appropriate to observe protein clustering at different stages. This feature was essential to capture the progressive changes in membrane curvature induced by clustering of VCC oligomers. The heterogeneity inherent to such liposomal systems also permitted us to observe the formation of different lipid-protein clusters and supramolecular assemblies in this study, which could otherwise be missed. Cryo-EM study of native cellular systems is difficult due to spillage of internal contents upon binding of pore-forming toxins, making particle-picking and classification challenging. Due to these aspects, liposomes serve as a great model membrane for studying protein-protein interaction on the membrane surface. (6, 9, 10, 13). The tendency of PFTs to aggregate upon liposomes even in short incubation period severely limited the application of liposomes in high-resolution structure determination of PFTs. Perforin, which is secreted by cytotoxic T lymphocytes or natural killer cells, has been documented to modulate the physicochemical properties of target membranes and aggregating upon lipid vesicles, causing invagination(14). Pneumolysin (9, 15–17), a Cholesterol-Dependent Cytolysin, has also been reported to rapidly aggregate upon liposomes(18, 19), limiting the toxin-membrane incubation period prior to cryo-ET data acquisition in one study(10). An Atomic Force Microscopy-based study of HlgAB with liposomes hypothesized that the target membrane blebbing was caused due to clustering of toxin oligomers and their proposed mechanism(20) was validated by our 2D class averages in this study, though on a different PFT-VCC. All of the abovementioned studies illustrate how membrane modulation and clustering by PFTs is a general phenomenon observed across different classes of PFTs.

Interestingly, we observe some similarities and differences in the behavior of VCC and HlgAB with respect to their interaction with the membrane. In our study with VCC, we observe the major population to be VCC oligomers bound to the membrane, with the minor population being larger superassemblies outside the membrane. Oligomers bound to the liposomes tended to laterally cluster together on the membrane. Unlike VCC, we saw HlgAB oligomers having a higher tendency to aggregate as per the cryo-EM micrographs. This could be possible due to the difference in membrane rupturing kinetics between VCC and HlgAB, since other factors (such as protein concentration, lipid environment, and incubation periods) were kept constant in the performed experiments. Additionally, VCC has a comparatively larger size (450 kDa) than HlgAB (270 kDa). During data processing, we were able to align the lipid membrane-embedded VCC, but not the smaller PFTs like HlgAB. Therefore, we were unable to identify toxin-induced membrane curvature in HlgAB.

Both of these toxins show the formation of larger superassemblies as a post-membrane damage state. We previously showed how HlgAB octamers formed an octahedral assembly with six octamers associated by their hydrophobic rim domain (6). However, in this study, we observed that VCC heptamers tended to adopt pentagonal pyramidal symmetry while forming larger superassemblies. In the case of HlgAB, we also observed the formation of these larger clusters at lower protein concentration and with erythrocytes, showing that the formation of these superassemblies was physiologically relevant(6). This also has important synthetic applications in the formation of nanocages since different oligomeric states of two PFTs have shown how octahedral and pentagonal pyramidal scaffolds can be obtained. Through grafting of viral virulence factors onto these highly symmetric nanocages, these can also possibly be used in vaccine development(21–24).

In this study, we focused on obtaining structural evidence and insights into protein clustering-induced membrane rupturing, however there are several interesting questions unanswered. We currently do not understand in depth the biophysical mechanism by which the clustering of toxins occurs in lipid membranes and how these oligomers induce blebbing. As a part of our future direction, we would like to obtain mechanistic insights into this phenomenon through complementary biochemical and molecular dynamics studies.

## CONCLUSIONS

Our current cryo-EM-based image analysis studies showed the PFTs, like VCC and γ-hemolysin, displayed blebbing during membrane rupture, resulting in the lipid-protein clusters. We also captured step-by-step membrane rupture incidents from our cryo-EM data, which demonstrated different membrane curvature. Additionally, multiple heptameric toxins deployed different tension on membranes, which led to various types of membrane curvature during the membrane rupture. This resulted as different types of protein-lipid clusters, where octameric toxins formed octahedral super assembly and heptameric toxins adopted pentagonal pyramidal architecture.

## METHODS

### Preparation of liposome-embedded VCC

Wild-type VCC was cloned, purified, and constituted into asolectin-cholesterol liposomes (1:1 wt/wt) as previously described(5). Briefly, cloning of the wild-type pro-VCC sequence was performed between the NdeI and BamHI restriction enzyme sites into the pET-14b vector (Novagen). Transformation was performed into Escherichia coli Origami B cells (Novagen) for expression and purification of recombinant protein. Purification of pro-VCC was done by Ni-nitrilotriacetic acid (QIAGEN) based affinity chromatography and Q-Sepharose anion-exchange chromatography (GE Healthcare). This was followed by proteolytic cleavage of the pro-domain using trypsin digestion in a protein:trypsin ratio of 2000:1 (wt/wt) for 5 min at 25 °C. Another round of Q-Sepharose anion-exchange chromatography. 6 μM mature VCC was then incubated with 1000 μg of asolectin:cholesterol liposomes (1:1 wt/wt) at 25°C for 1 hour. After treatment, ultracentrifugation was performed at 105,000 g for 30 min, following which the pellet was resuspended in 200 μl buffer containing 10 mM Tris-HCl (pH 8.0) and 150 mM NaCl and stored at −20°C until further use.

### Preparation of liposome-embedded γ-hemolysin

Wild-type γ-hemolysin subunits HlgA and HlgB were codon-optimized for cloning and expression in *E. coli* BL21 (DE3). Overexpression of both subunits was induced using 0.5 mM IPTG and incubated for 16 hours at 16°C. Two-step purification was performed for each protein using Ni-nitrilotriacetic acid (QIAGEN) based affinity chromatography and Superdex 200 increase 10/300 GL column (Cytiva) as previously described(6). Subunits HlgA and HlgB (in 1:1 molar ratio) were then treated with large unilamellar vesicles constituting equimolar concentrations of eggPC and cholesterol at 37°C for 30 minutes before proceeding for cryo-EM plunge-freezing.

### Cryo-EM data collection for proteo-liposomes and image processing for VCC

Quantifoil R1.2/1.3 300 mesh gold grids were glow discharged in a GloQube glow discharge system (Quorum Technologies Ltd.) for 130 s at 20 mA before sample addition. Then, 3 μl of liposome-embedded VCC/HlgAB was applied, incubated for 10 sec and excess sample was blotted for 5.5 s at 100% humidity before plunge freezing into liquid ethane using an FEI Vitrobot IV plunger. We collected ∼3500 multi-frame movies over 40 frames using 200 kV Talos Arctica TEM equipped with K2 Summit direct electron detector at magnification of 42000x and pixel size of 1.17 Å. In RELION3.1(25), Motion-corrected micrographs were used for manual particle picking and an initial 2D classification was performed to generate templates for further automatic particle picking, that yielded ∼259938 particles. Particles were extracted with a box size of 480 pixels and then imported into CryoSPARCv4.1(26). Rigorous and iterative 2D classifications were performed to yield 2D classes representing different states in liposomes and the superassemblies separately. Particles corresponding to superassemblies were used to generate an *ab initio* model and a homogeneous refinement was performed without any symmetry. Rigid body fitting was performed using UCSF ChimeraX(27).

## DATA AVAILABILITY STATEMENT

The data that support the findings of this study are available from the corresponding author upon request.

## AUTHOR CONTRIBUTIONS

SM and SD designed the experiments. SM performed the cryo-EM image processing and prepared the figures. SM and SD wrote the manuscript with inputs from KC. **CRediT statement: Suman Mishra:** Conceptualization; Data curation; Formal analysis; Visualization; Writing - original draft; Methodology; Investigation; Writing - review & editing; Validation. **Kausik Chattopadhyay:** Writing - review & editing. **Somnath Dutta:** Writing - review & editing; Funding acquisition; Conceptualization; Formal analysis; Visualization; Writing - original draft; Supervision; Project administration; Resources; Validation.

## ETHICAL STATEMENT

All experiments were performed in accordance with the guidelines laid by the Indian Institute of Science, Bangalore.

